# Contextual Fear Conditioning in Zebrafish

**DOI:** 10.1101/068833

**Authors:** Justin W. Kenney, Ian C. Scott, Sheena A. Josselyn, Paul W. Frankland

## Abstract

Zebrafish are a genetically tractable vertebrate that hold considerable promise for elucidating the molecular basis of behavior. Although numerous recent advances have been made in the ability to precisely manipulate the zebrafish genome, much less is known about many aspects learning and memory in adult fish. Here, we develop a contextual fear conditioning paradigm using an electric shock as the aversive stimulus. We find that contextual fear conditioning is modulated by shock intensity, prevented by inhibition of (N-methyl-D-aspartate) NMDA receptors, lasts at least 14 days, and exhibits extinction. Furthermore, fish of various background strains (AB, Tu, and TL) are able to acquire fear conditioning, but differ in fear extinction rates. Taken together, we find that contextual fear conditioning in zebrafish shares many similarities with the widely used contextual fear conditioning paradigm in rodents. Combined with the amenability of genetic manipulation in zebrafish, we anticipate that our paradigm will prove to be a useful complementary system in which to examine the molecular basis of vertebrate learning and memory.

## Introduction

Zebrafish are a useful model organism for studying physiology due to their genetic accessibility, low cost, and potential for high-throughput analysis (Grunwald and Eisen, 2002). Given that approximately 70% of human genes have an obvious orthologue in fish (Howe et al., 2013) zebrafish can be used to model various human diseases (Lieschke and Currie, 2007) and are emerging as a powerful tool for the *in vivo* screening of compounds for drug discovery (Zon and Peterson, 2005; MacRae and Peterson, 2015). More recently, both larval and adult zebrafish have been successfully utilized to study both basic questions in neuroscience, and gain a deeper understanding of the genetics of neuropsychiatric and neurodegenerative diseases (Agetsuma et al., 2010; Ahrens et al., 2012; Schmid and Haas, 2013; Kalueff et al., 2014; Leung and Mourrain, 2016).

The strength of zebrafish for understanding the genetic basis of behavior has been realized using forward genetic screens in both adult and larval fish (Darland and Dowling, 2001; Muto et al., 2005; Gerlai, 2010). With the advent of scalable genome editing technologies, such as clustered regularly interspaced short palindromic repeats (CRISPR)/Cas9, the potential for performing targeted high throughput reverse genetic screens is now also feasible (Hwang et al., 2013; Varshney et al., 2015). Additionally, zebrafish hold considerable potential in personalized medicine, as the CRISPR/Cas9 system has been successfully used to precisely knock-in exogenous DNA (Hisano et al., 2015). However, to fully leverage the zebrafish model to understand how genetics contributes to behavior in both health and disease requires a deep understanding of zebrafish behavior.

Adult zebrafish exhibit a rich repertoire of behaviors, from complex social interactions, to anxiety-like behaviors, and various forms of learning (Kalueff et al., 2013; Gerlai, 2015). Associative learning is an important, and highly conserved, form of learning in which an initially neutral conditioned stimulus (CS) is paired with an unconditioned stimulus (US) resulting in the expression of a conditioned response (CR) upon subsequent exposure to the CS. Although both classical (Pavlovian) and operant conditioning has been demonstrated in zebrafish (Arthur and Levin, 2001; Xu et al., 2007; Blank et al., 2009; Agetsuma et al., 2010; Sison and Gerlai, 2010; Valente et al., 2012; Manuel et al., 2014; Gorissen et al., 2015; Fernandes et al., 2016), many of these tasks require habituation or training over multiple days, do not last beyond 24 hours following training, are difficult to assess, or are not generalizable to fish of different genetic backgrounds. Here, we describe the development of a contextual fear conditioning task in adult zebrafish that is robust, rapidly acquired, straightforward to measure, and allows for the examination of the various phases of learning (acquisition, consolidation, and retrieval).

## Results

### Shock Intensity

We initially determined an appropriate shock intensity to reliably elicit both an unconditioned response (UR) and a CR in zebrafish. We administered shocks ranging from 0 to 20 mA to separate groups of fish during the training period (Figure 1A). We found that a 20 mA shock resulted in both a robust UR (i.e. an activity burst during shock administration) and CR (i.e. a decrease in distance travelled following repeated shock administration) whereas a 10 mA shock resulted in a less consistent UR and no CR, and the 5 mA shock resulted in no discernible change in locomotor activity (Figure 1B).

**Figure 1.**
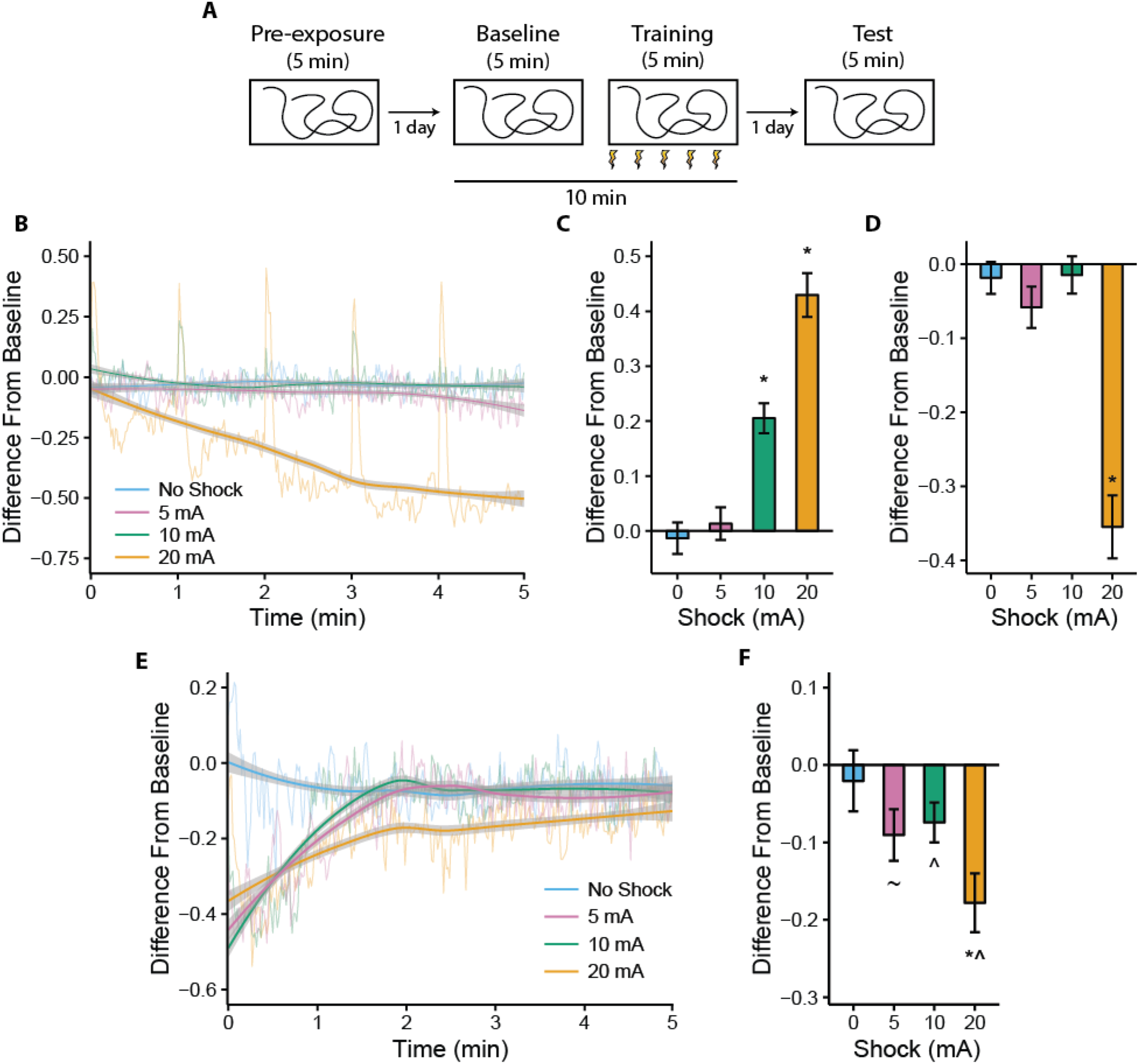
Contextual fear conditioning in adult zebrafish. **A**) Scheme indicating behavioral procedure used for inducing fear conditioning in zebrafish. **B**) Second-by-second difference scores during training in contextual fear conditioning. **C**) Difference scores at shock administration during training. **D**) Difference scores over the last 2.5 minutes of training. **E**) Second-by-second difference scores during the five minute test session. **F**) Difference scores over the first 2.5 minutes testing. Semi-transparent lines are average second-by-second data. Solid lines are the result of a local polynomial regression fit with 95% confidence interval for the fit (gray ribbons). ^*^-p < 0.05 compared to un-shocked fish, ∧ - p < 0.05 compared to a difference score of zero, ~ - p < 0.10 compared to a difference score of zero, n = 19-20.

The UR of fish to the shock stimulus differed depending on the shock intensity administered (F(3,74) = 41.8, P = 6.6 × 10^-16^) with a Dunnett post-hoc tests indicating that the 10 mA and 20 mA groups differed from the no shock group (5 mA: P = 0.88, 10 mA: P < 1.0 × 10^-4^, 20 mA: P < 1.0 × 10^-4^ Figure 1C). Additionally, the distance travelled of fish subjected to different shock intensities was different during the last half of the training trial (F(3,74) = 29, P = 1.6 × 10^-12^) with a Dunnett post-hoc test indicating that only the 20 mA group differing significantly from the no shock group (5 mA: P = 0.68, 10 mA: P = 0.99, 20 mA: P < 1.0 × 10^-4^; Figure 1D).

When placed back into the tank during the test, all three groups administered shocks initially had a decrease in their locomotor activity, however, the 5 and 10 mA groups returned to baseline levels of activity within 2 minutes, whereas the decrease in swimming in the 20 mA group was persistent throughout the trial (Figure 1E). Examination of the first half of the testing trial confirmed that there was a stimulus dependent effect of treatment on distance travelled (F(3,74) = 3.6, P = 0.018) with a Dunnett post-hoc test indicating that only the 20 mA group differed from the no shock group (5 mA: P = 0.35, 10 mA: P = 0.54, 20 mA: P = 0.0053; Figure 1F). However, Bonferroni corrected one-sample t-tests found a strong trend towards a difference from zero for the 5 mA group and that both the 10 and 20 mA groups significantly differed from zero (no shock: P = 1, 5 mA: P = 0.058, 10 mA: P = 0.038, 20 mA: P = 0.00018). Taken together, these data suggest that administration of 20 mA shocks results in the formation of a robust CS-US (tank-shock) association with less pronounced effects using the 5 and 10 mA shock intensities. Furthermore, the effect on behavior during testing appears to be most pronounced during the first half of the testing trial (Figure 1E). Based on these findings, we used the 20 mA shock intensity throughout the rest of our experiments and focus our analysis on the first 2.5 minutes of testing.

### Tank Specificity

The decrease in swimming we observe following shock administration to zebrafish may reflect injury, and not a conditioned fear response. Therefore, we sought to determine if fear conditioning in zebrafish is context-specific, a well-known characteristic of contextual fear conditioning in rodents (Owen et al., 1997; Rudy and O’Reilly, 1999). To accomplish this, we interspersed exposure to another novel tank (tank B) with exposure to the tank in which shocks were administered (tank A; Figure 2A). When fish were tested in tank A, they had a persistent decrease in their distance travelled relative to baseline compared to testing in tank B (Figure 2B). Comparing fish during the first half of the trial confirmed that testing in tank A resulted in a decrease in locomotor activity compared to testing in tank B (t(58) = 2.71, P = 0.0088; Figure 2C). These data suggest that zebrafish are able to discriminate between tanks, and thus the conditioned response is unlikely to be due to injury.

**Figure 2.**
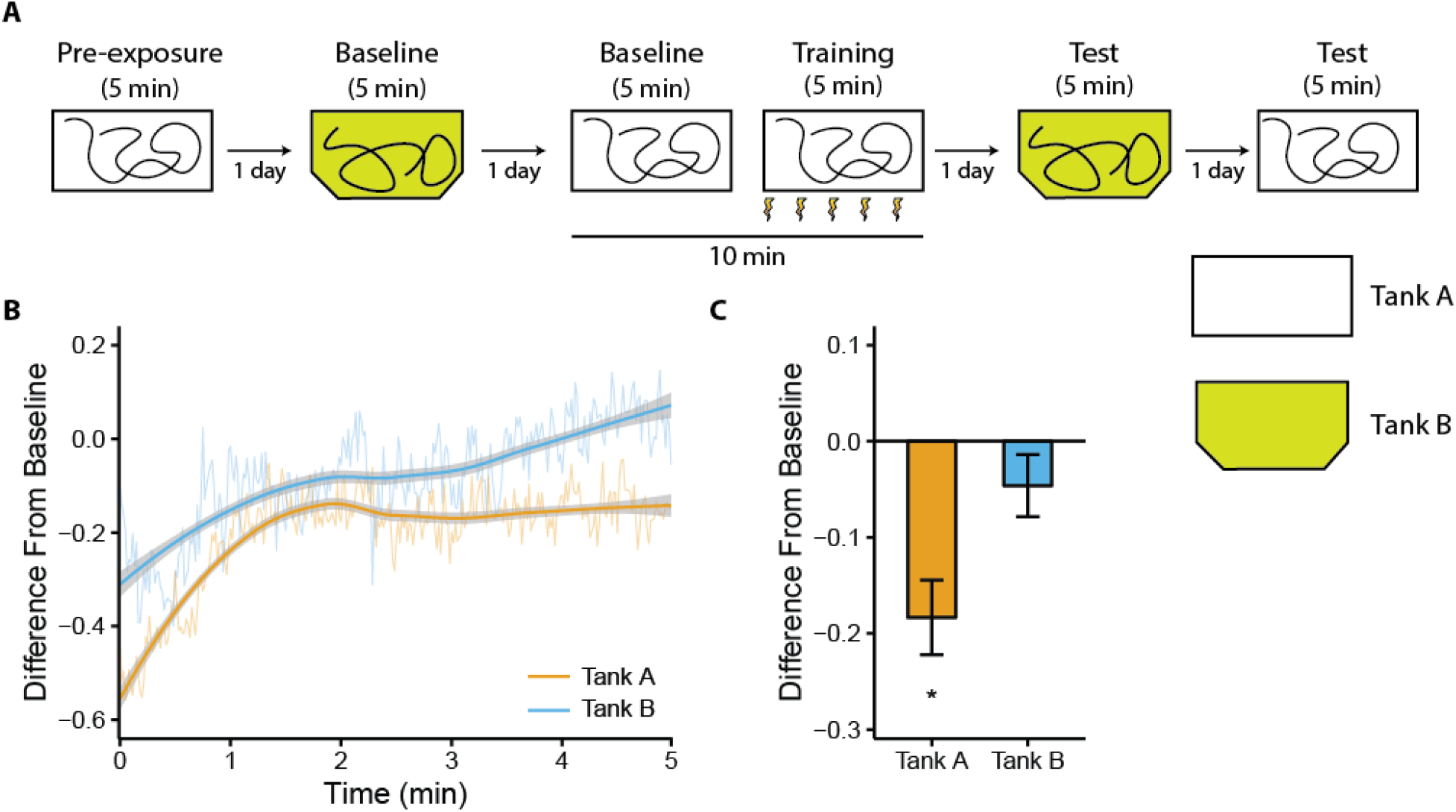
Contextual (tank) discrimination in adult zebrafish. **A**) Scheme indicating the behavioral procedure used to test for fear memory generalization. **B**) Second-by-second difference scores during testing in contextual fear conditioning in either tank A or tank B. **C**) Difference scores over the first 2.5 minutes of the test session. Semi-transparent lines are average second-by-second data. Solid lines are the result of a local polynomial regression fit with 95% confidence interval for the fit (gray ribbons). ^*^ * - p < 0.05 compared to testing in Tank B, n = 30.

### MK-801

Many forms of learning in a variety of species are known to depend on NMDA receptor function (Abel and Lattal, 2001), including zebrafish (Blank et al., 2009; Sison and Gerlai, 2011). To determine if contextual fear conditioning similarly requires NMDA function, we administered 20 μM MK-801, an NMDA receptor antagonist, to zebrafish following training in contextual fear conditioning (Figure 3). We found that fish treated with vehicle had a persistent decrease in locomotor activity during testing, whereas those treated with MK-801 did not (Figure 3A). An examination of the first half of testing confirmed that vehicle treated fish suppressed their locomotor activity compared to MK-801 treated fish (t(35) = 3.45, P = 0.0015; Figure 3B).

**Figure 3.**
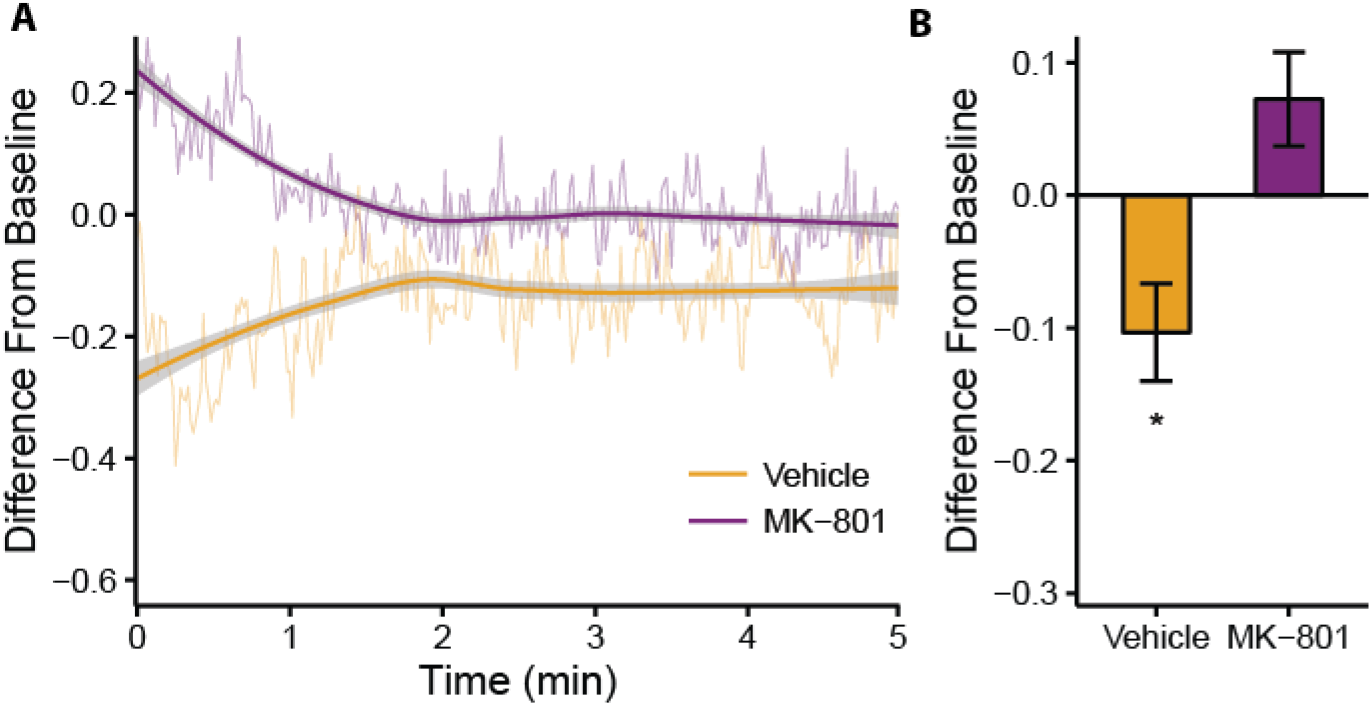
The effect of MK-801 administration after training on contextual fear conditioning. **A**) Second-by-second difference scores during testing in contextual fear conditioning in fish administered vehicle or MK-801 (20 μM). **B**) Difference scores over the first 2.5 minutes of testing of fear conditioning. Semi-transparent lines are average second-by-second data. Solid lines are the result of a local polynomial regression fit with 95% confidence interval for the fit (gray ribbons). ^*^- p < 0.05 compared to MK-801 treated fish, n = 18-19.

### Time Course

Thus far we have found that contextual fear memories are present one day following training, which is a time frame consistent with numerous previous studies examining memory in zebrafish (Blaser and Vira, 2014; Kalueff et al., 2014; Gerlai, 2015) However, it is largely unknown how long memories in zebrafish may last. In order to determine how long-lasting the contextual fear conditioning memory is, we trained separate groups of zebrafish with delays of 7, 14, 21, or 28 days between training and testing (Figure 4). We found that at all delays, the decrease in locomotor activity at the beginning of testing was clearly present (Figures 4A-D, left). Analysis of the first half of the testing trials indicated that a robust fear memory lasts between 14 and 21 days (7 days: t(36) = 3.25, P = 0.0025; 14 days: t(38) = 2.80, P = 0.0079; 21 days: t(37) = 1.89, P = 0.066; 28 days: t(39) = 1.53, P = 0.13; Figures 4A-D, right).

**Figure 4.**
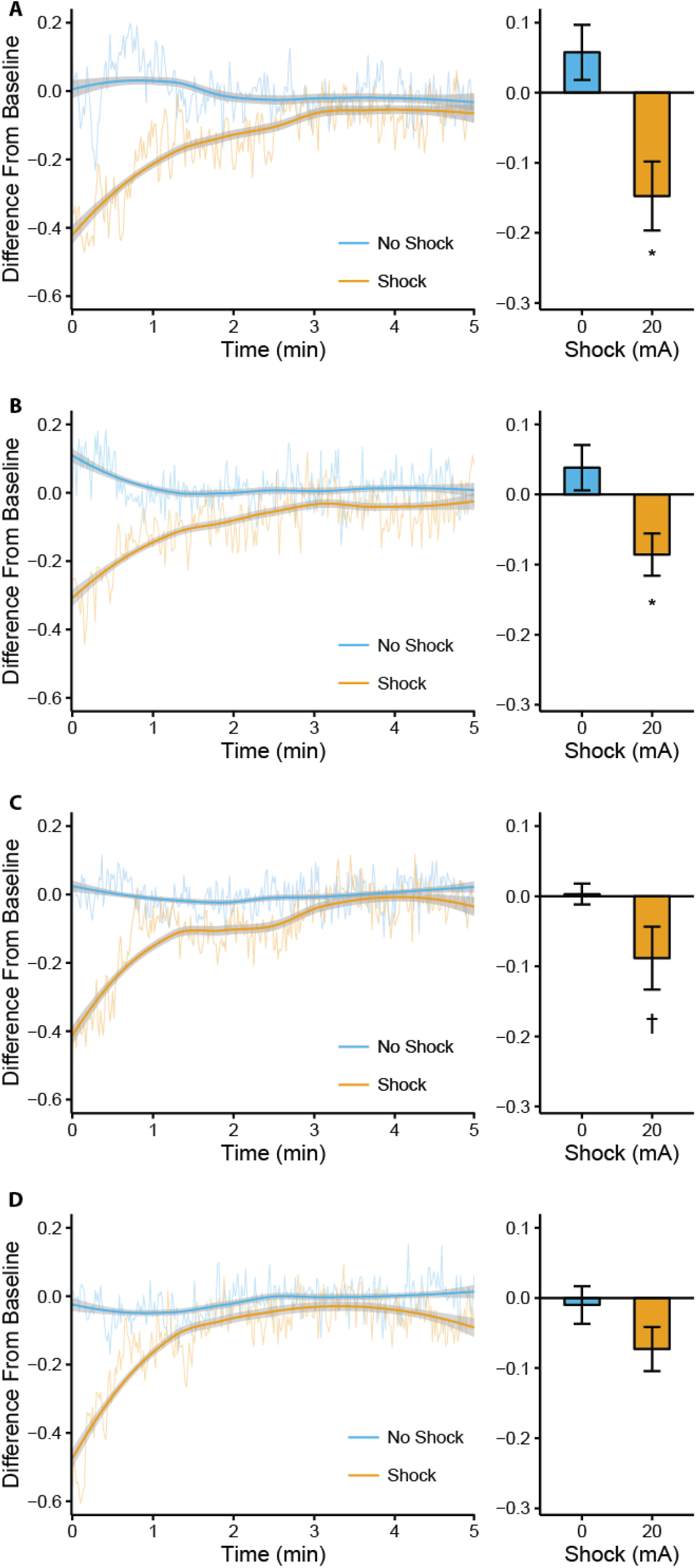
Effect of different retention delays on contextual fear conditioning in adult zebrafish. Fish were tested after a delay of 7 (**A**), 14 (**B**), 21 (**C**), or 28 (**D**) days following training. Second-by-second difference scores during the entire test (left) and during the first 2.5 minutes of testing (right). Semi-transparent lines are average second-by-second data. Solid lines are the result of a local polynomial regression fit with 95% confidence interval for the fit (gray ribbons). ^*^ - p < 0.05, † - p < 0.10 compared to un-shocked fish, n = 19-21.

### Extinction

One phenomenon related to associative learning is extinction. In extinction, repeated exposure to the CS in the absence of the US results in a reduction of the CR. We attempted to extinguish the tank-shock association by repeated exposure of the fish to the conditioning tank on successive days. We found that, although within session extinction is present during each of the four days of testing, the decrease in locomotor activity at the beginning of the session persisted throughout the days of testing (Figure 5A). A 2 × 4 (group × day) repeated measures ANOVA applied to the change in distance travelled during the first half of each testing trial found a main effect of group (F(1,61) = 31.5, P = 5.13 × 10^-7^), but not day (F(3,183) = 2.11, P = 0.10), and a day by group interaction (F(3,183) = 4.44, P = 0.0049; Figure 5B). Bonferroni corrected post-hoc tests found that shocked fish differed from un-shocked fish at each day (day 1: P = 1.68 × 10^-7^, day 2: P = 0.0019, day 3: P = 0.026, day 4: P = 0.026), and that the shocked fish at days 3 and 4, but not day 2, differed from shocked fish at day 1 (compared to day 1, day 2: P = 0.073, day 3: P = 0.0038, day 4: P = 0.0034). Taken together, these data suggest that the memory partially extinguished from day 1, but was still present after four consecutive days of exposure to the conditioning tank.

**Figure 5.**
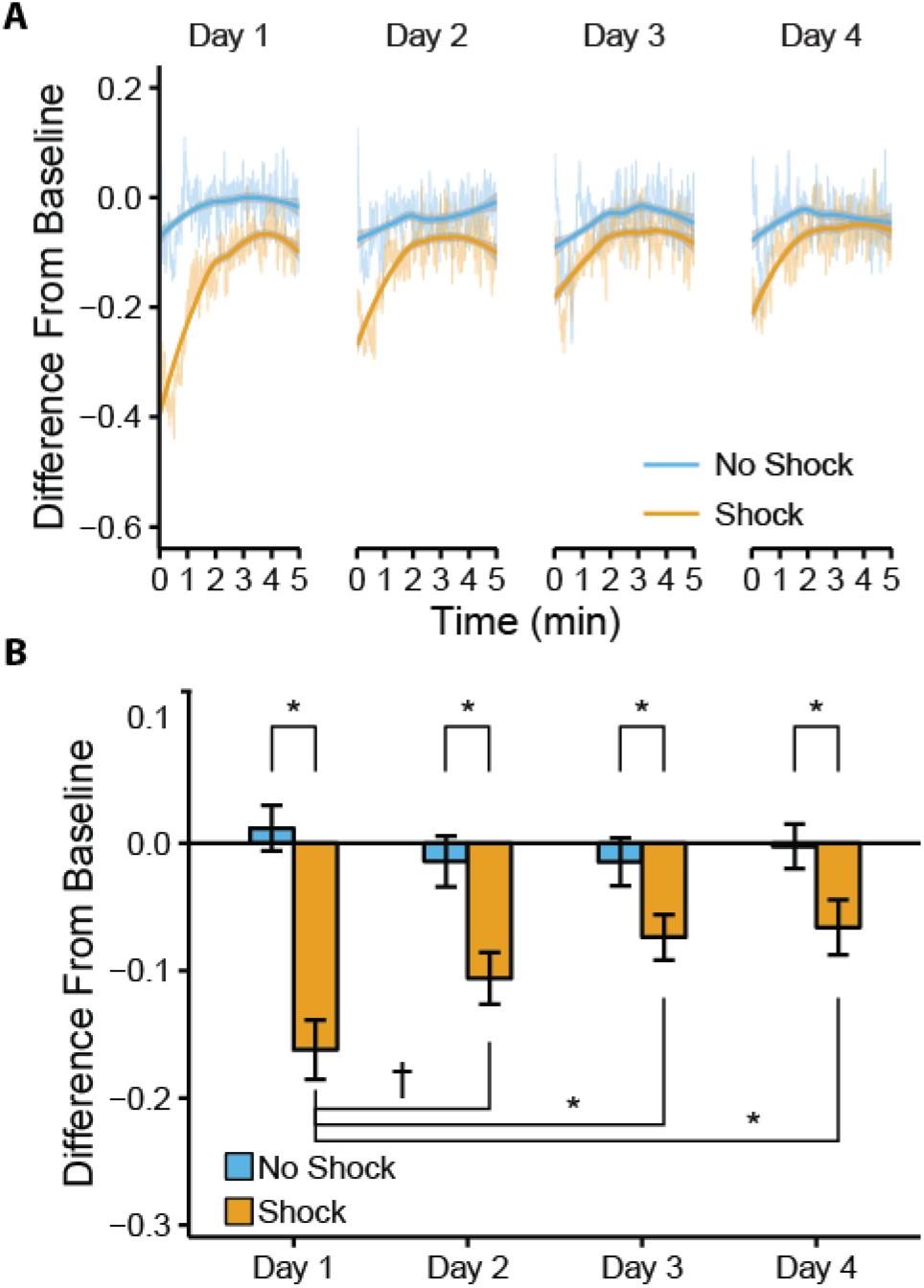
Extinction of contextual fear conditioning in adult zebrafish. **A**) Second-by-second difference scores during each of 4 successive days of testing following training in contextual fear conditioning. **B**) Difference scores over the first 2.5 minutes of 4 successive days of testing following training in contextual fear conditioning. Semi-transparent lines are average second-by-second data. Solid lines are the result of a local polynomial regression fit with 95% confidence interval for the fit (gray ribbons). ^*^ - p < 0.05, † - p < 0.10, n = 31-32.

### Tu and TL fish

Many learning tasks in rodents, and some in fish, are known to be affected by the genetic background of the strain under study (Owen et al., 1997; Gerlai, 1998; Vignet et al., 2013; Gorissen et al., 2015). Thus, we sought to determine if different widely used strains of zebrafish exhibit responses in our contextual fear conditioning paradigm that are similar to the AB strain we have examined thus far. Tu fish have been used for sequencing of the zebrafish genome (Howe et al., 2013) and TL fish are another commonly used strain that are genetically distinct from both the AB and Tu strains (Trevarrow and Robison, 2004; Guryev et al., 2006). Fish of both the Tu and TL background exhibited responses during training that were similar to that of AB fish, although the decrease in swimming in Tu fish appeared to plateau more quickly than in either the AB or TL strains (Figures 6A and B, left). Both Tu and TL fish increased their locomotor activity in response to the 20 mA shock (Tu: t(30) = 4.13, P = 0.0027; TL: t(36) = 3.30, P = 0.0022). An examination of data during successive exposures to the testing tank indicated that Tu fish had both robust within and between session extinction (Figure 6C) whereas TL fish showed little within session extinction and only moderate between session extinction (Figure 6D).

**Figure 6.**
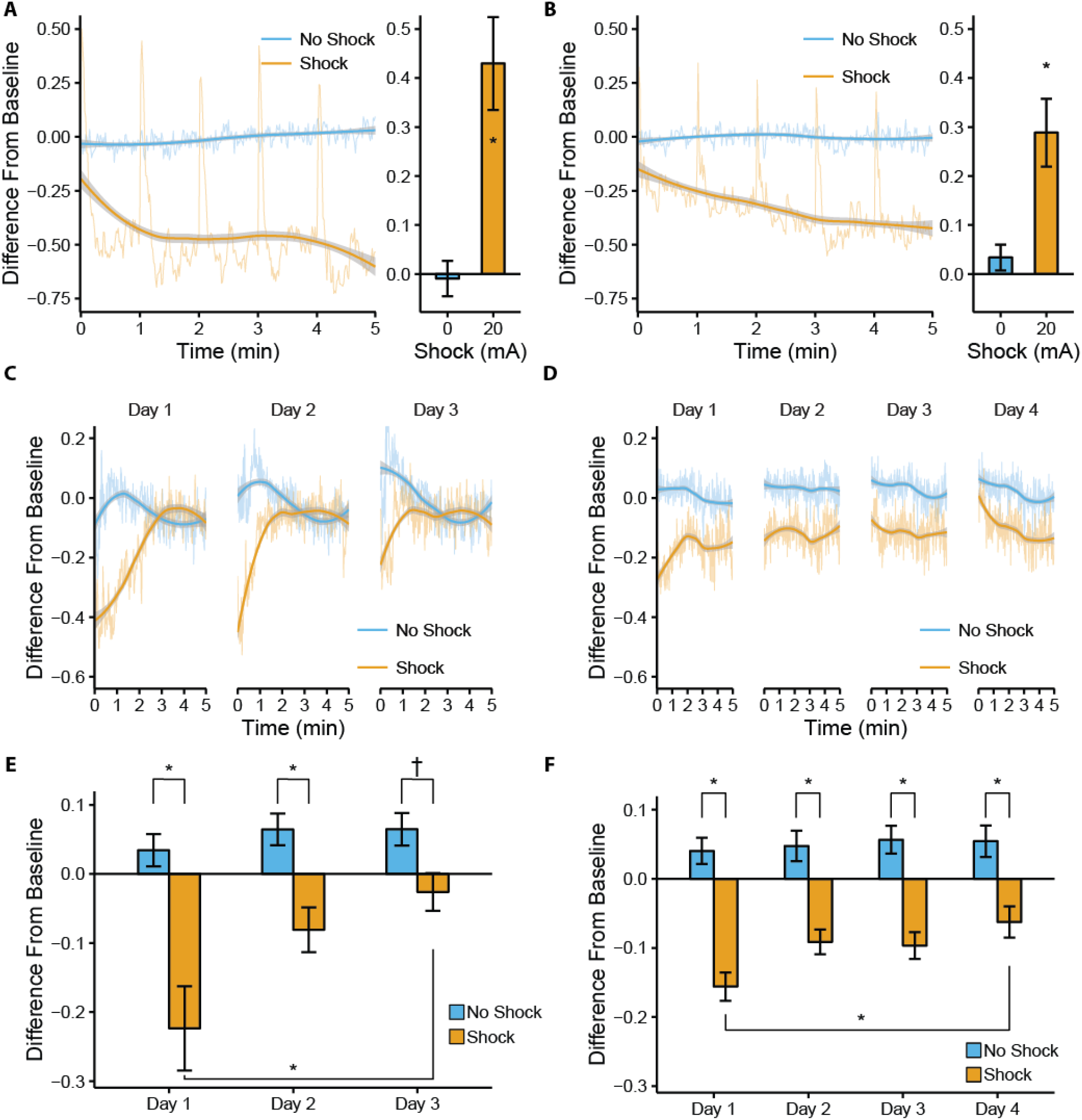
Contextual fear conditioning in Tu and TL zebrafish strains. Behavior during training in Tu (**A**) and TL (**B**) fish with second-by-second data on the left and response to the shock on the right. Second-by-second difference scores during successive days of testing in Tu (**C**) and TL (**D**) fish. Difference scores during the first 2.5 minutes of 3 or 4 successive days of testing in Tu (**E**) and TL (**F**) fish. Semi-transparent lines are average second-by-second data. Solid lines are the result of a local polynomial regression fit with 95% confidence interval for the fit (gray ribbons). ^*^- p < 0.05, † - p < 0.10, n = 15-20.

In Tu fish, a 2 × 3 (group × day) repeated measures ANOVA performed on the difference scores during the first half of testing during each extinction day found a main effect of group (F(1, 30) = 28.0, P = 1.0 × 10^-5^), a main effect of day (F(2, 60) = 6.44, P = 0.0029) and a strong trend towards an interaction between day and group (F(2,60) = 3.02, P = 0.056; Figure 6E). Bonferroni corrected post-hoc tests indicated that shocked Tu fish differed from un-shocked fish at days 1 and 2, with a trend towards a difference on day 3 of extinction (day1: P = 0.0038, day 2: P = 0.005, day 3: P = 0.087). Furthermore, comparisons with the first day of testing indicated that fish significantly decreased their activity on day 3, but not day 2, of extinction testing (compared to day 1, day 2: P = 0.24, day 3: P = 0.037).

In TL fish, a 2 × 4 (group × day) repeated measures ANOVA using difference score data from the first half of testing on each day found a main effect of group (F(1, 36) = 79.8, P = 1.16 × 10^-10^), and a main effect of day (F(3,108) = 2.97, P = 0.035), but no group by day interaction (F(3, 108) = 1.50, P = 0.22; Figure 6F). Bonferroni corrected post-hoc tests comparing shocked and un-shocked fish for each day found that shocked fish differed from un-shocked fish at each day (day 1: P = 2.19 × 10^-7^, day 2: P = 0.00016, day 3: P = 2.4 × 10^-5^, day 4: P = 0.0056) and that in shocked fish only testing on day 4 was different from the first day of testing (compared to day 1, day 2: P = 0.16, day 3: P = 0.30, day 4: P = 0.028). Taken together, these data suggest that extinction takes longer and is less robust in TL fish compared to both AB and Tu fish.

## Discussion

In the present study, we find that contextual fear conditioning in zebrafish is a robust and rapidly-acquired task that allows for the examination of numerous types of learning. Contextual fear conditioning can be modulated by the strength of the shock, lasts at least 14 days, extinguishes following repeated exposure to the context (tank), and can be acquired by fish with a variety of genetic backgrounds. Furthermore, the apparatus was built from inexpensive off-the-shelf components and the analyses were performed using open-source software, thereby making this behavioral paradigm easily scalable and widely accessible.

The simplest alternative explanation for the findings in the present study is that the decrease in locomotor activity observed following shock administration is due to injury and not the formation of a tank-shock association. However, several aspects of our findings argue against this interpretation. Firstly, in almost every experiment, there is significant within session extinction where the decrease in locomotor activity is greatest during the first half of the test session and approaches zero during the second half of testing. If the fish were injured by the shock, we would expect the decrease in locomotor activity to remain constant throughout the session. Secondly, if fish were injured due to the shock, it is unlikely that administration of MK-801 would be able to prevent learning in this paradigm (Figure 3). Thirdly, fish are able to modulate their decrease in locomotor activity based on the tank in which they are placed where the decrease is greater in the tank they were shocked in as opposed to a different novel tank (Figure 2). If the suppression of locomotor activity were due to injury, we would expect the decrease to be the same in both tanks. Finally, we see no effect on locomotor activity after placement back in home tanks immediately following shock administration when any such effect of injury would be most obvious (unpublished observations).

Although a number of other aversive learning tasks have been described in zebrafish, none have been reported to be as rapidly acquired, robust, and generalizable as the contextual fear conditioning task we describe here. For example, inhibitory avoidance in zebrafish has been found to be rapidly acquired, requiring only one trial for learning (Blank et al., 2009). However, some strains of fish, such as the widely used AB strain, are unable to learn inhibitory avoidance (Gorissen et al., 2015) whereas we find that the AB, Tu, and TL strains all demonstrate robust learning of contextual fear conditioning. Both operant and classical fear conditioning to a discrete visual cue have been described in adult zebrafish using a shock (Agetsuma et al., 2010; Valente et al., 2012) or alarm pheremone as the US (Hall and Suboski, 1995). However, work using shocks required multiple trials with the memory lasting no longer than 6 hours (Agetsuma et al., 2010; Valente et al., 2012), and the alarm pheromone only elicits an unconditioned response in ~40% of fish (Hall and Suboski, 1995). In contrast, we find that our contextual fear conditioning memory lasts at least 14 days (Figure 4) and the shock US elicits a measurable, robust, and consistent increase in locomotor activity across several strains of fish (Figures 1 and 6).

To the best of our knowledge, the present study is the first demonstration of memory retention beyond 24 hours after one-trial learning in zebrafish. Establishment of a learning task with a clearly delineated acquisition phase and sufficient strength to last weeks is useful as it allows for the study of not only molecular/cellular consolidation that occurs over 24 hours following learning, but also slower processes (e.g, systems consolidation or forgetting) that may emerge over the course of days and weeks rather than hours (Abel and Lattal, 2001; Wixted, 2004; Frankland and Bontempi, 2005). Interestingly, we found that in zebrafish the strength of the fear memory decreased progressively as the interval between training and testing increased (Figure 4). This is in contrast to what is observed in rodents, where fear memories tend to have the same or greater strength at increasing delays (Fanselow et al., 1994; Houston et al., 1999).

In addition to contextual fear learning, our learning paradigm can also be used to study extinction. Extinction of associative memories has previously been reported in zebrafish for inhibitory avoidance (Piato et al., 2011), visual discrimination learning (Colwill et al., 2005), and conditioned place avoidance (Yu et al., 2006). Here, we find that extinction occurs both within a single testing session as well as between sessions during repeated exposure to the testing tank. Interestingly, we find that fish of the Tu background extinguish after three days of exposure to the test tank whereas fish from the AB and TL genetic backgrounds still have significantly elevated levels of fear even after four days of extinction. This is similar to what is observed in the extinction of contextual fear in rodents, where complete extinction of the fear response occurs in some strains of rodents but not others (Stiedl et al., 1999; Camp et al., 2009).

Taken together, we find contextual fear conditioning in adult zebrafish to be a reliable, long-lasting, and versatile task for studying various aspects of associative learning. The paradigm described here is similar in many ways to contextual fear conditioning in rodents that has proven to be a powerful tool for probing the many facets of learning and memory (Johansen et al., 2011; Maren et al., 2013). The robustness of fear conditioning, along with the ability to modulate the strength of learning via shock intensity, makes it suitable to examine factors that may either enhance or inhibit learning.

## Materials and Methods

### Subjects

Subjects were AB, Tu, or TL Zebrafish 3-6 months of age. Male and female fish were housed together, 7-11 per 2 L tank. All fish were bred and raised at the Hospital for Sick Children in high density racks under standard conditions with a 14:10 light/dark cycle (lights on at 8:30). Fish were fed twice daily with *Artemia salina*. Behavioral testing took place between 12:00 and 18:00. All procedures were approved by the Hospital for Sick Children Animal Care and Use Committee.

### Apparatus

The behavioral apparatus consisted of two identical Aquaneering crossing tanks (ZHCT100T) covered with white self-adhesive film and two horizontal black stripes to provide visual cues. Two stainless steel mesh grids were also placed at each end of the tank. Tanks were surrounded by white Plasticore to prevent interference from external visual stimuli. Webcams (Logitech C270) were mounted approximately 35 cm above tanks. An alternate tank (Tank B) consisted of a 1.8 L Aquaneering tank (ZT180) with blue and green stripes on the bottom. To further differentiate the alternate tank from the training tank, several visual stimuli (photographs of Norwegian Fjords) were placed on the inside walls of the Plasticore enclosure. A Nextech (c-215) camera was placed approximately 40 cm in front of the tank. Videos were recorded on a laptop PC using Free Screencast software at approximately 15 frames per second and encoded using the Xvid MPEG-4 codec. DC electric shocks were administered using a Bio-Rad PowerPac Basic power supply.

### Behavioral Procedure

The basic behavioral procedure had three phases (**Figure 1A**): pre-exposure, baseline/training, and testing. During pre-expsoure on day 1, fish were placed individually in the test tank for five minutes. On day 2, baseline and training, individual fish were placed in the test tank for 10 minutes. Locomotor activity during the first half of the session was considered baseline. During the second half of the session, fish received five shocks (3 seconds, 5-20 mA) spaced one minute apart starting 5 minutes into the session. On day 3, individual fish were placed back in the test tank for five minutes. Video was recorded for each session and saved for offline analysis. Before and after each individual fish was placed in the apparatus, tanks were rinsed with distilled water and filled with 800 mL of fresh system water. Clear plastic lids were placed on the top of the tank during recording. At least two separate tanks of fish were used for each experimental group.

### Behavioral Analysis

Videos of individual fish were tracked using Ctrax (Branson et al., 2009) and the distance travelled during each second was calculated using a custom written Python script (available at http://github.com/jkenney9a/Zebrafish). A difference score was used to quantify the change in swimming activity during a given test period compared to the baseline period:

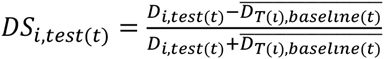

Where DS is the difference score, D is distance travelled, *i* is an individual fish, *t* is time, and T(*i*) is the tank that fish *i* is from. Because the fish were group housed and we could not reliably distinguish individual fish from one another, the average distance travelled for a given tank was used as the baseline measure (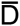). When calculating the difference score during training, the following equation was used:

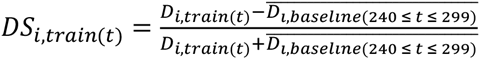

Where the baseline distance travelled was calculated as the average distance the individual fish travelled during each second of the last minute of baseline.

Data from training and testing in fear conditioning are presented in two formats: 1) Average difference scores for each second of training or testing calculated as described above. Given the considerable noise in second by second data, these data are fit using a local polynomial regression to provide a measure of the overall trend in the data. 2) An average of difference scores during the first or second half (2.5 minutes) of training or testing. We use data generated from this method for inferential statistics to determine group differences and are presented as the mean ± SEM.

### Drug Administration

A stock solution of MK-801 (0924; Tocris) was diluted into 100 mL of facility water. Fish were placed in a 250 mL semi-opaque beaker with MK-801 (20 μM) or vehicle (water) immediately following training in fear conditioning for 15 minutes.

### Statistical Analyses

Data were analyzed with one-way ANOVAs and two-way repeated measures ANOVAs as appropriate. Dunnett post-hoc tests were used to compare multiple groups to un-shocked fish as indicated. Experiments with two groups were analyzed using independent samples t-tests. When multiple pair-wise comparisons were made using one-sample or independent samples t-tests, results were corrected using the Bonferroni method as indicated. All statistical analyses was performed using R (v3.1.3).

## Acknowledgments

We would like to thank Angela Morley, Alan Ng, and Monica Yu for excellent care of zebrafish and facility maintenance. We would also like to thank Robert Gerlai, Yohaan Fernandes, Laura McDonald, Sarah Hutchinson, and Emanuela Pannia for helpful discussions. JWK was supported by a Human Frontiers Science Program long-term fellowship (LT000759/2014). This work was supported by a grant from the Canadian Institute for Health Research to PWF (FDN143227).

## References

Abel T, Lattal KM (2001) Molecular mechanisms of memory acquisition, consolidation and retrieval. Curr Opin Neurobiol 11:180–187.

Agetsuma M, Aizawa H, Aoki T, Nakayama R, Takahoko M, Goto M, Sassa T, Amo R, Shiraki T, Kawakami K, Hosoya T, Higashijima S, Okamoto H (2010) The habenula is crucial for experience-dependent modification of fear responses in zebrafish. Nat Neurosci 13:1354–1356.

Ahrens MB, Li JM, Orger MB, Robson DN, Schier AF, Engert F, Portugues R (2012) Brain-wide neuronal dynamics during motor adaptation in zebrafish. Nature 485:471.

Aizenberg M, Schuman EM (2011) Cerebellar-Dependent Learning in Larval Zebrafish. J Neurosci 31:8708–8712.

Arthur D, Levin E (2001) Spatial and non-spatial visual discrimination learning in zebrafish (Danio rerio). Anim Cogn 4:125–131.

Blank M, Guerim LD, Cordeiro RF, Vianna MRMM (2009) A one-trial inhibitory avoidance task to zebrafish: Rapid acquisition of an NMDA-dependent long-term memory. Neurobiol Learn Mem 92:529–534.

Blaser RE, Vira DG (2014) Experiments on learning in zebrafish (Danio rerio): A promising model of neurocognitive function. Neurosci Biobehav Rev 42:224–231.

Branson K, Robie AA, Bender J, Perona P, Dickinson MH (2009) High-throughput ethomics in large groups of Drosophila. Nat Methods 6:451–457.

Camp M, Norcross M, Whittle N, Feyder M, D’Hanis W, Yilmazer-Hanke D, Singewald N, Holmes A (2009) Impaired Pavlovian fear extinction is a common phenotype across genetic lineages of the 129 inbred mouse strain. Genes, Brain Behav 8:744–752.

Colwill RM, Raymond MP, Ferreira L, Escudero H (2005) Visual discrimination learning in zebrafish (Danio rerio). Behav Processes 70:19–31.

Darland T, Dowling JE (2001) Behavioral screening for cocaine sensitivity in mutagenized zebrafish. Proc Natl Acad Sci 98:11691–11696.

Fanselow MS, Kim JJ, Yipp J, De Oca B (1994) Differential Effects of the Af-Methyl-D-Aspartate Antagonist DL-2-Amino-5-Phosphonovalerate on Acquisition of Fear of Auditory and Contextual Cues. Behav Neurosci 108:235–240.

Fernandes YM, Rampersad M, Luchiari AC, Gerlai R (2016) Associative learning in the multichamber tank: A new learning paradigm for zebrafish. Behav Brain Res 312:279–284.

Frankland PW, Bontempi B (2005) The organization of recent and remote memories. Nat Rev Neurosci 6:119–130.

Gerlai R (1998) Contextual learning and cue association in fear conditioning in mice: a strain comparison and a lesion study. Behav Brain Res 95:191–203.

Gerlai R (2010) High-Throughput Behavioral Screens: the First Step towards Finding Genes Involved in Vertebrate Brain Function Using Zebrafish. Molecules 15:2609–2622.

Gerlai R (2015) Zebrafish phenomics: Behavioral screens and phenotyping of mutagenized fish. Curr Opin Behav Sci 2:21–27.

Gorissen M, Manuel R, Pelgrim TNM, Mes W, de Wolf MJS, Zethof J, Flik G, van den Bos R (2015) Differences in inhibitory avoidance, cortisol and brain gene expression in TL and AB zebrafish. Genes, Brain Behav 14:428–438.

Grunwald DJ, Eisen JS (2002) Headwaters of the zebrafish — emergence of a new model vertebrate. Nat Rev Genet 3:717–724.

Guryev V, Koudijs MJ, Berezikov E, Johnson SL, Plasterk RHA, Eeden FJM van, Cuppen E (2006) Genetic variation in the zebrafish. Genome Res 16:491–497.

Hall D, Suboski MD (1995) Visual and Olfactory Stimuli in Learned Release of Alarm Reactions by Zebra Danio Fish (Brachydanio rerio). Neurobiol Learn Mem 63:229–240.

Hisano Y et al. (2015) Precise in-frame integration of exogenous DNA mediated by CRISPR/Cas9 system in zebrafish. Sci Rep 5:8841.

Houston FP, Stevenson GD, McNaughton BL, Barnes CA (1999) Effects of Age on the Generalization and Incubation of Memory in the F344 Rat. Learn Mem 6:111–119.

Howe K et al. (2013) The zebrafish reference genome sequence and its relationship to the human genome. Nature 496:498–503.

Hwang WY, Fu Y, Reyon D, Maeder ML, Tsai SQ, Sander JD, Peterson RT, Yeh J-RJ, Joung JK (2013) Efficient genome editing in zebrafish using a CRISPR-Cas system. Nat Biotechnol 31:227–229.

Johansen JP, Cain CK, Ostroff LE, LeDoux JE (2011) Molecular mechanisms of fear learning and memory. Cell 147:509–524.

Kalueff A V., Stewart AM, Gerlai R (2014) Zebrafish as an emerging model for studying complex brain disorders. Trends Pharmacol Sci 35:63–75.

Kalueff A V et al. (2013) Towards a comprehensive catalog of zebrafish behavior 1.0 and beyond. Zebrafish 10:70–86.

Leung LC, Mourrain P (2016) Drug discovery: Zebrafish uncover novel antipsychotics. Nat Chem Biol 12:468–469.

Lieschke GJ, Currie PD (2007) Animal models of human disease: zebrafish swim into view. Nat Rev Genet 8:353–367.

MacRae CA, Peterson RT (2015) Zebrafish as tools for drug discovery. Nat Rev Drug Discov 14:721–731.

Manuel R, Gorissen M, Piza Roca C, Zethof J, Vis H van de, Flik G, Bos R Van Den (2014) Inhibitory Avoidance Learning in Zebrafish (Danio Rerio): Effects of Shock Intensity and Unraveling Differences in Task Performance. Zebrafish 11:341–352.

Maren S, Phan KL, Liberzon I (2013) The contextual brain: implications for fear conditioning, extinction and psychopathology. Nat Rev Neurosci 14:417–428.

Muto A, Orger MB, Wehman AM, Smear MC, Kay JN, Page-McCaw PS, Gahtan E, Xiao T, Nevin LM, Gosse NJ, Staub W, Finger-Baier K, Baier H (2005) Forward Genetic Analysis of Visual Behavior in Zebrafish. PLoS Genet 1:e66.

Owen EH, Logue SF, Rasmussen DL, Wehner JM (1997) Assessment of learning by the Morris water task and fear conditioning in inbred mouse strains and F1 hybrids: implications of genetic background for single gene mutations and quantitative trait loci analyses. Neuroscience 80:1087–1099.

Piato ÂL, Capiotti KM, Tamborski AR, Oses JP, Barcellos LJG, Bogo MR, Lara DR, Vianna MR, Bonan CD (2011) Unpredictable chronic stress model in zebrafish (Danio rerio): Behavioral and physiological responses. Prog Neuro-Psychopharmacology Biol Psychiatry 35:561–567.

Rudy JW, O^’^Reilly RC (1999) Contextual fear conditioning, conjunctive representations, pattern completion, and the hippocampus. Behav Neurosci 113:867–880.

Schmid B, Haas C (2013) Zebrafish as an animal model for neurodegeneration. J Neurochem 127:461–470.

Sison M, Gerlai R (2010) Associative learning in zebrafish (Danio rerio) in the plus maze. Behav Brain Res 207:99–104.

Sison M, Gerlai R (2011) Associative learning performance is impaired in zebrafish (Danio rerio) by the NMDA-R antagonist MK-801. Neurobiol Learn Mem 96:230–237.

Stiedl O, Radulovic J, Lohmann R, Birkenfeld K, Palve M, Kammermeier J, Sananbenesi F, Spiess J (1999) Strain and substrain differences in context- and tone-dependent fear conditioning of inbred mice. Behav Brain Res 104:1–12.

Trevarrow B, Robison B (2004) Genetic Backgrounds, Standard Lines, and Husbandry of Zebrafish. In: Methods in Cell Biology, pp 599–616.

Valente A, Huang K-H, Portugues R, Engert F (2012) Ontogeny of classical and operant learning behaviors in zebrafish. Learn Mem 19:170–177.

Varshney GK, Pei W, LaFave MC, Idol J, Xu L, Gallardo V, Carrington B, Bishop K, Jones M, Li M, Harper U, Huang SC, Prakash A, Chen W, Sood R, Ledin J, Burgess SM (2015) High-throughput gene targeting and phenotyping in zebrafish using CRISPR/Cas9. Genome Res 25:1030–1042.

Vignet C, Bégout M-L, Péan S, Lyphout L, Leguay D, Cousin X (2013) Systematic Screening of Behavioral Responses in Two Zebrafish Strains. Zebrafish 10:365–375.

Wixted JT (2004) The Psychology and Neuroscience of Forgetting. Annu Rev Psychol 55:235–269.

Xu X, Scott-Scheiern T, Kempker L, Simons K (2007) Active avoidance conditioning in zebrafish (Danio rerio). Neurobiol Learn Mem 87:72–77.

Yu L, Tucci V, Kishi S, Zhdanova I V. (2006) Cognitive Aging in Zebrafish Miall C, ed. PLoS One 1:e14.

Zon LI, Peterson RT (2005) In vivo drug discovery in the zebrafish. Nat Rev Drug Discov 4:35–44.

